# Deciphering Immune Complexity: Single-Cell Insights into Autoimmune Myocarditis Progression

**DOI:** 10.1101/2024.03.12.584698

**Authors:** Farag Mamdouh, Waleed K. Abdulsahib, Dalal Sulaiman Alshaya, Eman Fayad, Refaat A. Eid, Ghadi Alsharif, Mohammed A. Alshehri, Hassan M. Otifi, Mohamed A. Soltan, Muhammad Alaa Eldeen

## Abstract

Autoimmune myocarditis is a complex inflammatory response in the heart caused by abnormal immune system activity. We used modern single-cell technologies to analyze the complex gene expression patterns in autoimmune myocarditis tissue samples during several stages of inflammation: acute, subacute, and chronic. We identified the presence of T cell-monocyte complexes in both control and myocarditis samples from different phases using detailed analysis of vast single-cell RNA sequencing data. These complexes were notably more prevalent throughout the acute and subacute stages. Our investigation of gene ontology revealed their involvement in important processes as signal transduction, immune response control, and T cell proliferation and activation. We conducted a thorough analysis of trajectories, uncovering the step-by-step changes of macrophages into clusters linked to antigen presentation, oxidative stress responses, complement activation, and phagocytosis. Furthermore, our examination of neutrophil paths revealed their development and maturation from the bone marrow to distinct functional stages. Our analysis of cellular communication networks revealed consistent and phase-specific patterns, providing valuable information on how immune responses change as the disease progresses. We noted a substantial increase in BAFF signaling from normal to acute phases, while CHEMERIN signaling was upregulated from acute to chronic phases. The discoveries greatly improve our understanding of autoimmune myocarditis on a molecular level, providing a strong basis for creating personalized precision medicine treatments for each patient.

**Graphical Abstract:** 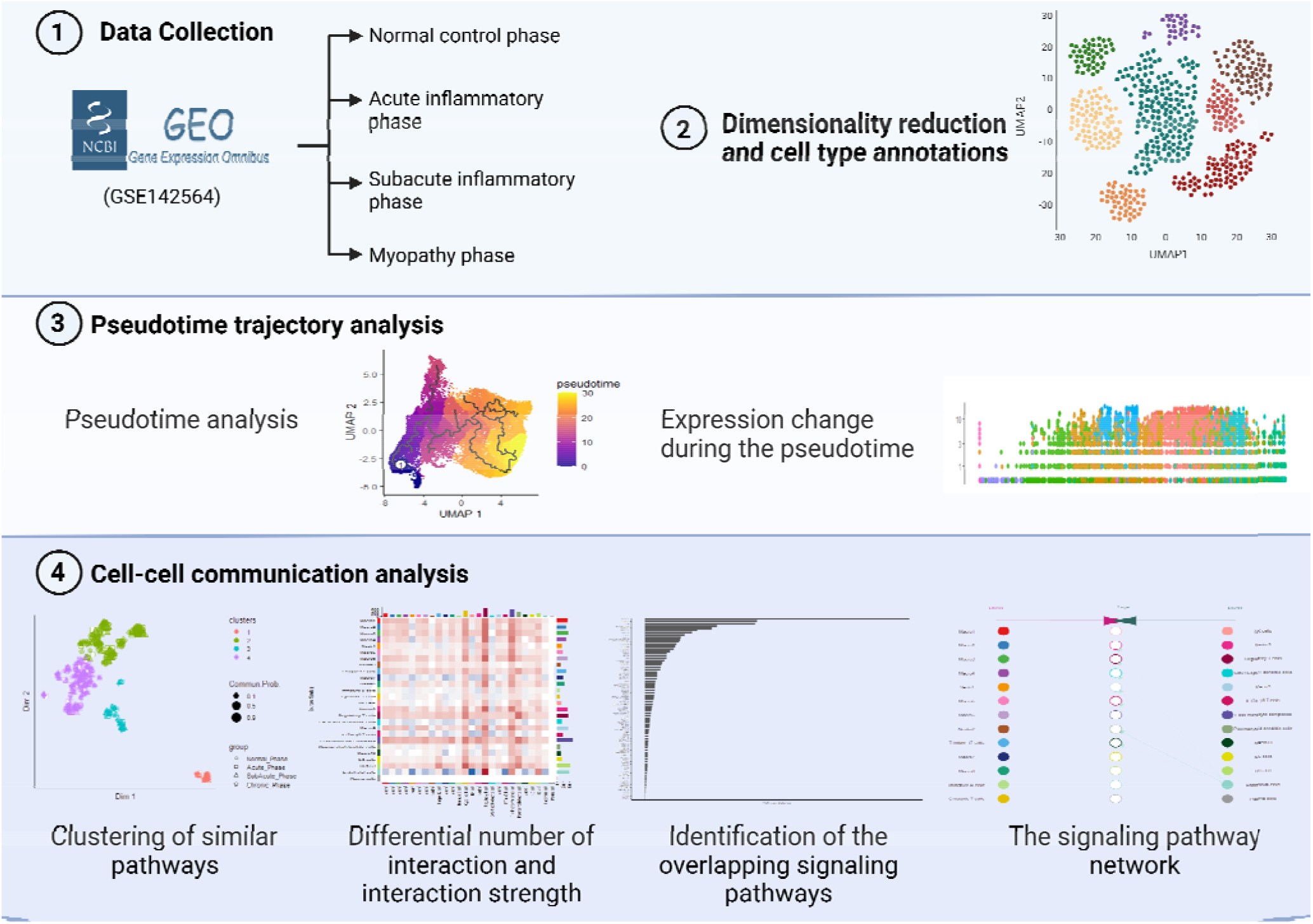

## 1. Introduction

Autoimmune diseases, caused by abnormalities in immunological tolerance systems, provide a significant problem in modern medicine. Disruptions in key components, including energy, control, and clonal deletion, play a role in their development [1,2]. Distinguished from autoimmune disorders, autoinflammatory conditions manifest as uncontrolled and nonspecific inflammatory responses, adding further complexity to the immunological landscape [3].

Myocarditis is a complex inflammatory condition involving inflammation of the heart muscle that advances through acute, subacute, and chronic phases [4]. Considerable progress has been achieved in understanding the fundamental processes of myocarditis, but the details of how immune cells enter cardiac tissue are still not comprehensively understood [5]. In response to this knowledge gap, the imperative arises to develop targeted interventions that selectively modulate autoimmunity or attenuate immune cell-mediated processes implicated in the development of inflammatory cardiomyopathy, all while minimizing collateral damage to the host.

Single-cell RNA sequencing (scRNA-seq) is an advanced method that integrates molecular biology and computational biology to get distinct understanding of the complex immunopathogenesis of autoimmune myocarditis [6]. Single-cell RNA sequencing surpasses the constraints of bulk transcriptomic methods by analyzing the transcriptional patterns of individual cells. This technique provides an in-depth comprehension of cellular variety, reveals new phenotypic characteristics, and exposes the changing cellular activity in the microenvironment [6]. The objective of our work is to understand the intricate immunological patterns that exist in autoimmune myocarditis. Our objective is to examine samples from various phases of inflammation and myopathy in order to get insight into the progression of the disease throughout time. Our approach entails examining the genetic behavior of cells, comprehending intercellular communication, and elucidating the process of inferring pseudo-time trajectories. This will aid in the identification of novel approaches for accurate medicinal interventions for autoimmune myocarditis.

## 2. Results

### 2.1. Characterization of Immune Cell Types across the Four Phases of Autoimmune Myocarditis

This study utilized a total of 42 single-cell RNA samples, comprising 10 control samples, 10 samples from the acute inflammatory phase, 9 samples from the subacute inflammatory phase, and 13 samples from the chronic myopathy phase. Access to the data is available in the Gene Expression Omnibus (GEO) database under accession number [GSE142564]. Distinct transcriptional patterns of various cell populations were discerned through dimensionality reduction and unsupervised cell clustering techniques employing the Seurat package [7]. Cell clusters were identified by analyzing their gene expression profiles, emphasizing important marker genes. The clusters were then classified into 26 primary cell types (Fig. 1A).

**Fig. 1.**
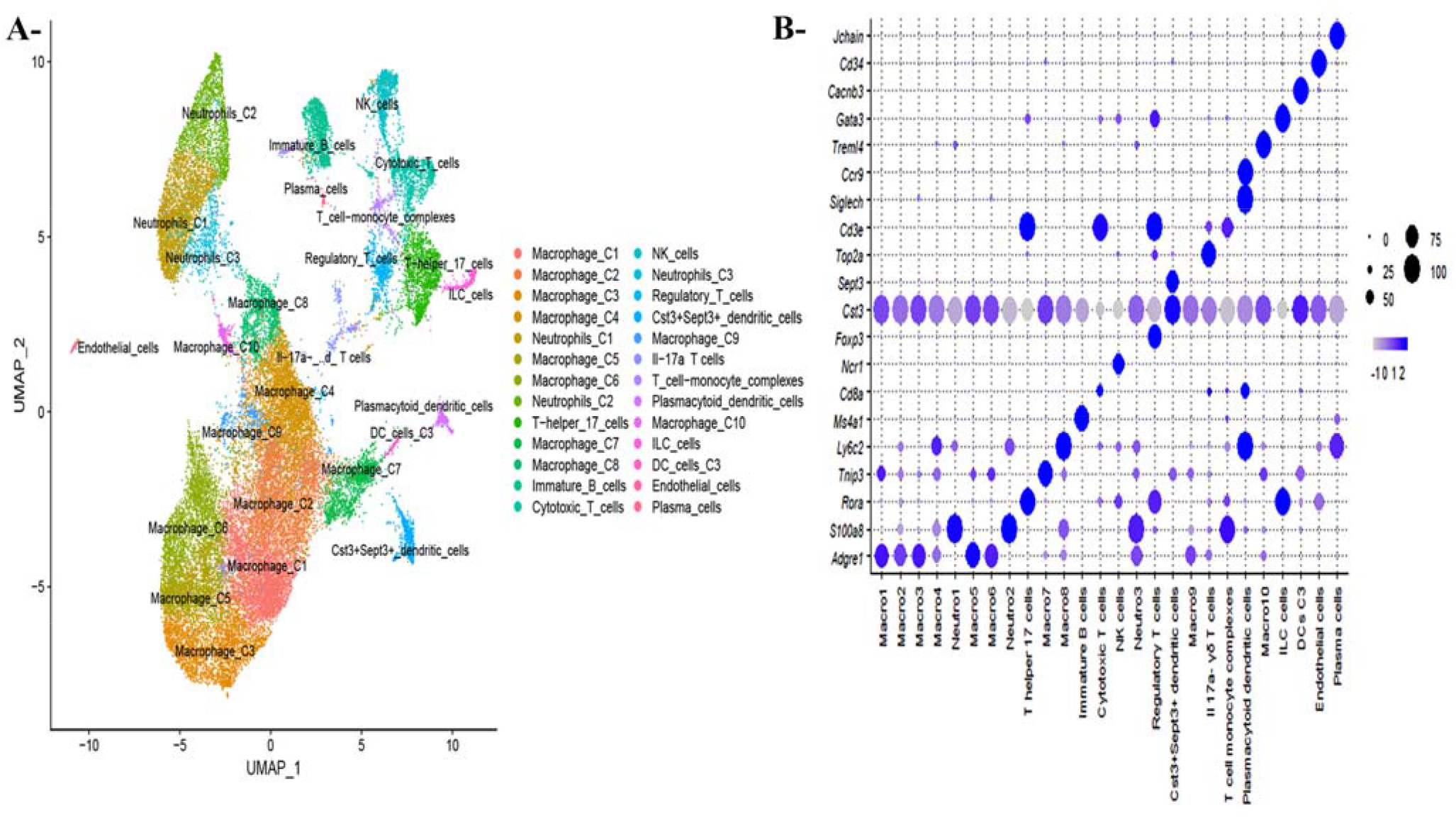
Transcriptomic Landscape and Heterogeneity of Cell Populations. (A) Uniform Manifold Approximation and Projection (UMAP) visualization illustrating the annotation of all cell types identified through single-cell RNA sequencing (scRNA-seq); (B) Barplot depicting the markers associated with each annotated cell type.

The identified major cell types encompass diverse macrophage subtypes implicated in antigen presentation, complement activation, phagocytosis, and response to oxidative stress, as elucidated in a recent publication [8]. Additionally, T cell subtypes (including T helper cells, cytotoxic T cells, regulatory T cells, Il17a^-γδ^ T cells, and T cell-monocyte complex cells), neutrophils, B cell subsets (immature B cells and plasma cells), dendritic cell types (cst^+^sept^+^ cluster and plasmacytoid dendritic clusters), innate lymphoid cells (ILCs), endothelial cells, and NK cells were identified.

### 2.2. Involvement of T Cell-Monocyte Complexes in the Progression of Autoimmune Myocarditis

Herein, We have discovered 315 T cell-monocyte complexes, highlighting the importance of intercellular communication in immune responses. Monocytes, which can differentiate into specialized antigen-presenting cells (APCs), including macrophages or dendritic cells (DCs)[9], are being acknowledged more for their involvement in adaptive immune responses [10,11]. Communication occurs either through direct cell-to-cell interaction or by the release of signaling molecules such as cytokines. Physical contacts between T cells and monocytes play a crucial role in initiating immune responses. Our analysis revealed that T cell-monocyte complexes were most abundant during the subacute and acute phases, decreasing during the chronic period and being least present during the normal phase (Fig. 2A). Furthermore, the expression of Cd4/Cd14 was significantly elevated in the T cell-monocyte complex [12,13] (Fig 2B).

**Fig. 2.**
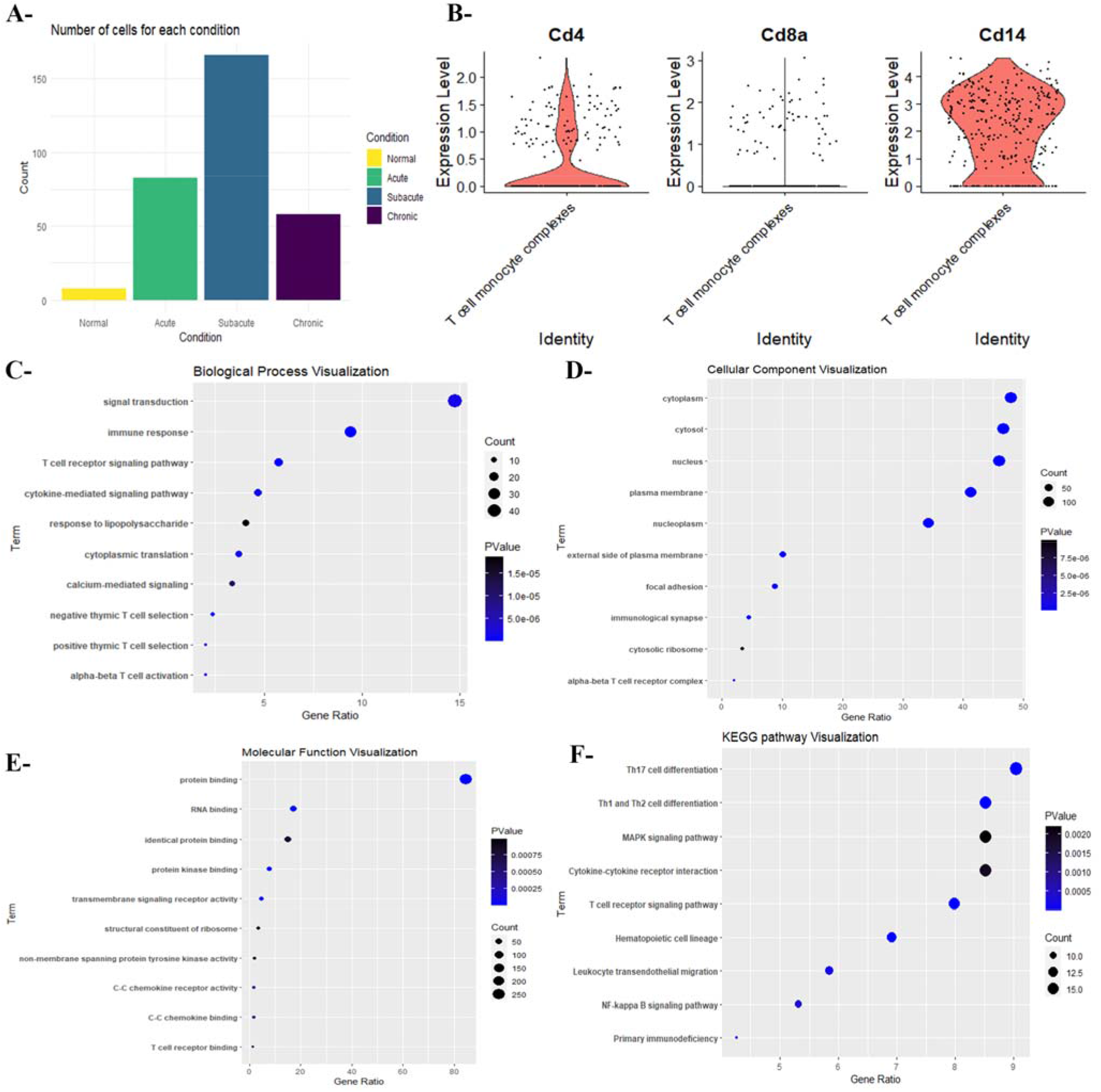
Characteristics of T Cell-Monocyte Complexes. (A) Bar plot illustrating the distribution of T cell-monocyte complexes across different phases; (B) Expression profile of marker genes associated with T cell-monocyte complexes; (C) Analysis of biological processes associated with differentially expressed genes (DEGs) in T cell-monocyte complexes; (D) Examination of cellular components represented by DEGs within T cell-monocyte complexes; (E) Evaluation of molecular functions attributed to DEGs within T cell-monocyte complexes; (F) KEGG pathway analysis highlighting the involvement of DEGs in critical pathways related to T cell-monocyte complex function.

An in-depth biological process study of differentially expressed genes (DEGs) in these complexes revealed their participation in signal transduction, immunological regulation, and T cell proliferation and activation pathways (Fig 2C). Our analysis of cellular components emphasized the significance of cytoplasmic, cytosolic, nuclear, and membrane structures for facilitating strong adhesion between T cells and monocytes (Fig. 2D). Our study delved deeper into molecular function to reveal the numerous activities of DEGs in T cell-monocyte complexes, such as protein binding, RNA binding, protein kinase binding, and participation in different signaling cascades (Fig. 2E).

KEGG pathway analysis underscored the substantial contribution of these complexes to critical pathways such as hematopoietic cell lineage establishment, T cell types differentiation, T cell receptor, and MAPK signaling, NF-kappa B Signaling Pathway, and Cytokine-Cytokine Receptor Interaction (Fig. 2F). These findings emphasize their crucial role in orchestrating adaptive immune responses, regulating inflammation, and facilitating cellular communication. In summary, our study offers a thorough investigation of the dynamic properties of T cell-monocyte complexes, emphasizing their crucial involvement in T cell growth, activation, and immunological responses.

### 2.3. Heterogeneity and Temporal Dynamics of Macrophages

Single-cell RNA sequencing (scRNA-seq) analysis revealed that macrophages constitute the predominant immune cell population within the heart. To further elucidate macrophage diversity, we performed a sub-clustering analysis, identifying 10 distinct sub-clusters (Fig. 3A). Notably, clusters Macro4 and Macro8 exhibited significant expression of the monocyte marker, suggesting their origin from monocytes that later differentiated into macrophages. Additionally, Macro5 showed elevated expression of Lyve1, F13a1, Cbr2, Cd163, Folr2, and Timd4 genes, associated with tissue-resident macrophages (supplementary Fig. 1) [14-17].

**Fig. 3.**
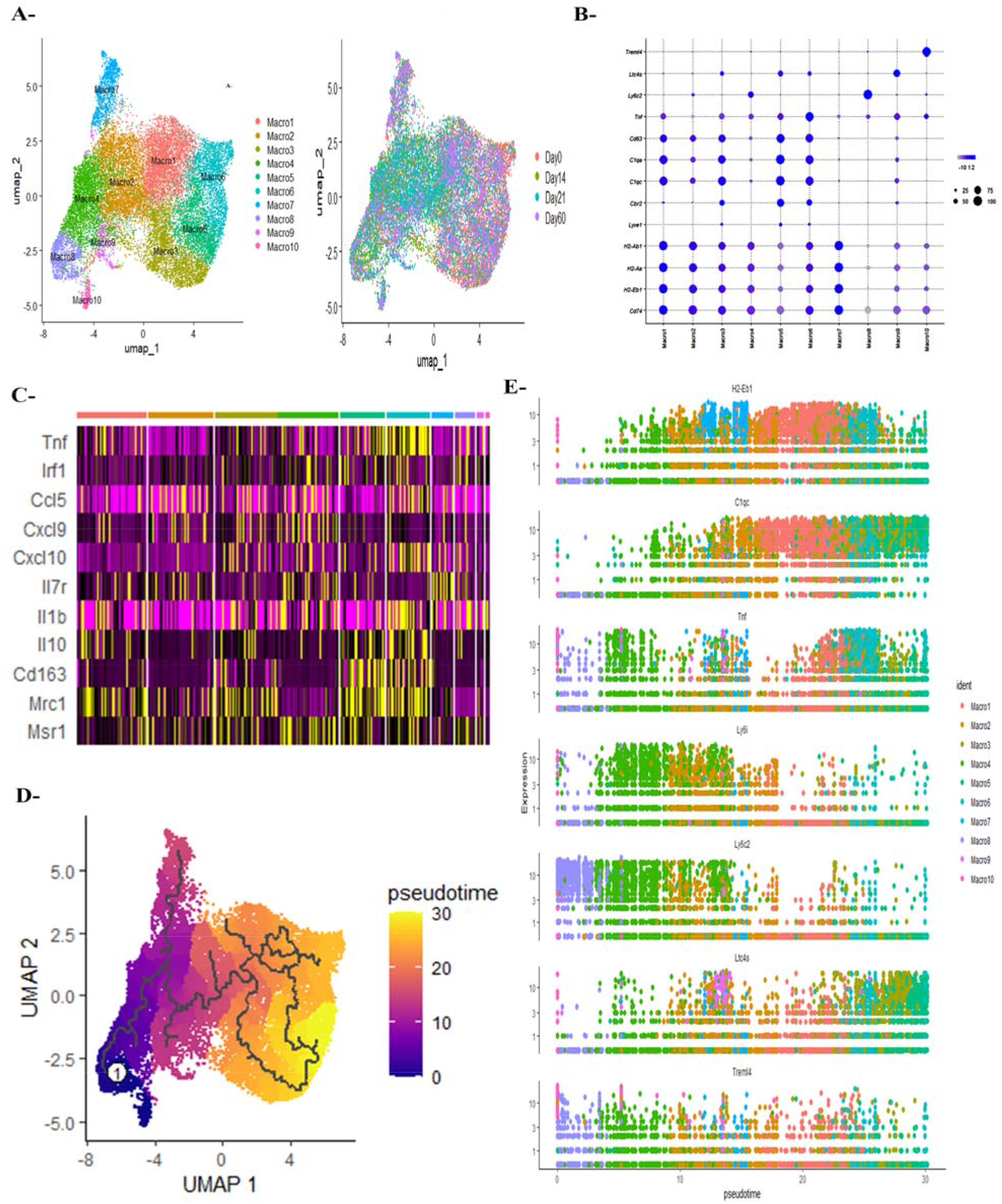
Developmental Trajectory of Macrophages in Autoimmune Myocarditis. (A) Uniform Manifold Approximation and Projection (UMAP) plot depicting the clustering of macrophage subtypes; (B) Dot plot illustrating the expression of marker genes for different macrophage types; (C) Heatmap displaying markers associated with M1 and M2 macrophages; (D) Pseudo-time trajectory analysis depicting the developmental trajectory of macrophages; (E) Expression changes of marker genes during pseudo-time advancement, highlighting dynamic gene expression patterns throughout macrophage development.

Macrophage clusters Macro1, Macro2, and Macro7 showed increased expression of H2-Eb1, H2-Aa, H2-Ab1, and Cd74 that is associated with antigen presentation, suggesting they are tissue-resident Mhc-II macrophages [17]. On the other hand, Macro3, Macro5, and Macro6 showed elevated gene expression related to the Complement and Phagocytosis processes [8]. Furthermore, Macro9 had high levels of Ltc4s expression, suggesting oxidative stress, while Macro10 had elevated Treml4 expression, linked to increased inflammatory activities (Fig. 3b) [8,18]. However, we did not identify subgroups accurately representing M1- or M2-polarized macrophages (Fig. 3C). We analyzed the course of monocyte/macrophage infiltration into damaged cardiac areas using the Monocle R package [19] to understand the underlying mechanisms. Cells originated from Ly6c2^hi^ monocytes and followed a trajectory, shifting from macrophage-derived monocytes to clusters related to antigen presentation and oxidative stress pathways, then progressing to pathways involving Complement and Phagocytosis (Fig. 3D). Expression profiles of marker genes within each cell group were assessed over pseudo-times, revealing dynamic changes (Fig. 3E). Notably, expression levels of Ly6i and Ly6c2 markers in macrophages decreased over pseudo-time, while genes associated with antigen presentation, complement, and phagocytosis clusters showed sustained or increased expression. H2-Eb1 exhibited a drop towards the end or pseudo-time, but C1qc showed a sustained increase. Ltc4s, indicating oxidative stress, exhibited consistent upregulation, whereas Tnf and Treml4 displayed regular expression patterns, except oxidative stress macrophage pseudo-time (Fig. 3E).

### 2.4. Heterogeneity and Temporal Dynamics of Neutrophils

Neutrophils play a crucial role in the immune response by migrating from the bloodstream to infected tissues in response to inflammatory signals, regulating inflammation in the cardiovascular system [20-22]. Despite their significance in various inflammatory conditions, their role in myocarditis has been largely overlooked [23]. Here, we hypothesized that neutrophils contribute significantly to the initial immunological response and are responsible for the impairment of heart function.

Utilizing UMAP clustering, we identified three distinct neutrophil clusters (Fig. 4A). Interestingly, the normal group exhibited a lower number of neutrophils, while the acute and subacute phases showed a higher abundance of neutrophils (Fig. 4B). The Neutro3 cluster showed elevated levels of Tuba1b and Rpl12, indicating its bone marrow origin and the differentiation of neutrophils, consistent with earlier research [24]. Neutro2 was associated with neutrophil degranulation, leukocyte migration, and actin cytoskeleton organization, while Neutro3 showed involvement in myeloid leukocyte migration, inflammatory response, and neutrophil degranulation [25]. Additionally, the Neutro1 cluster showed elevated levels of Oasl2 and Il18bp expression, suggesting its involvement in regulating defensive responses, neutrophil degranulation, and cytokine production.

**Fig. 4.**
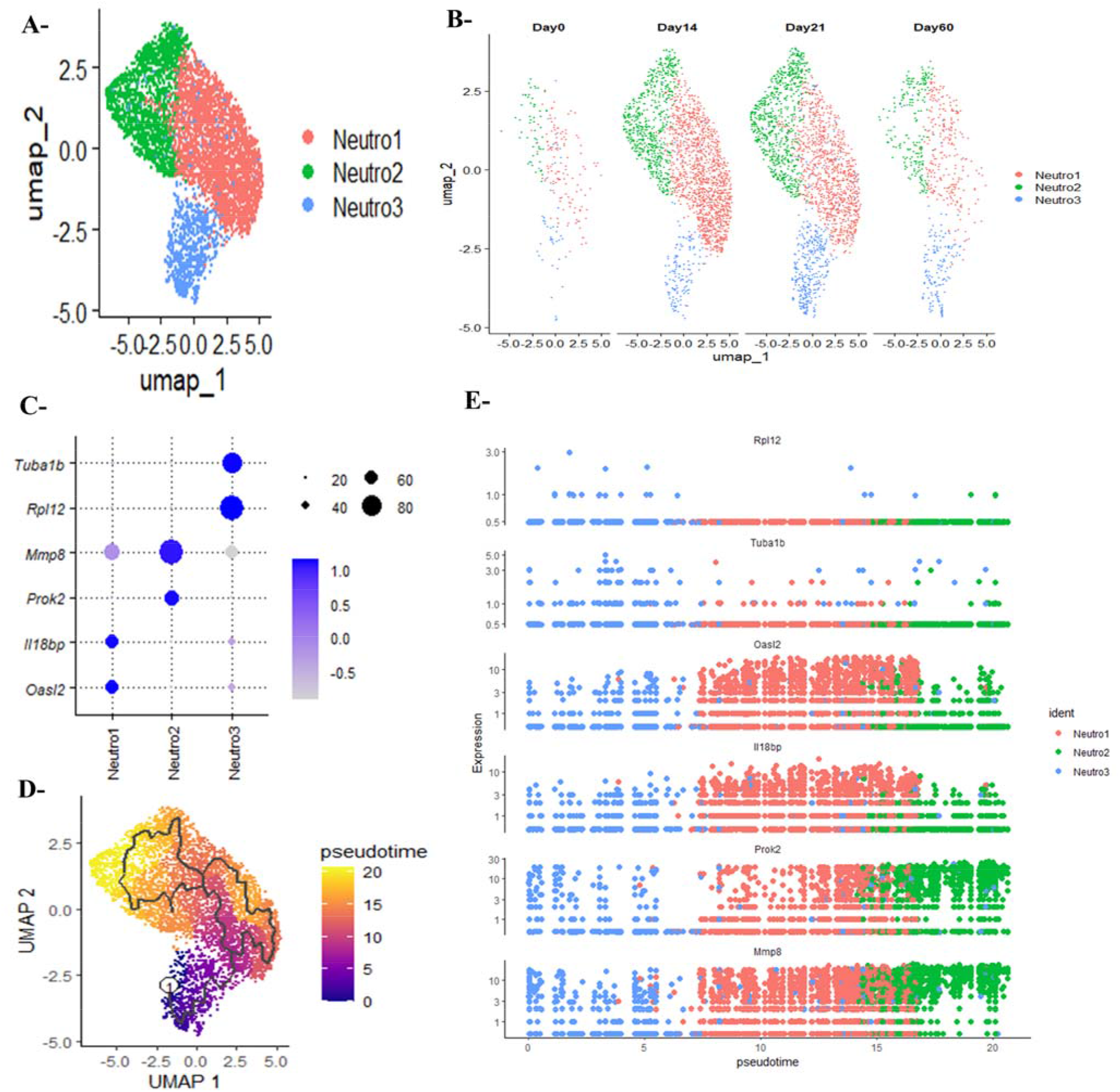
Developmental Trajectory of Neutrophils in Autoimmune Myocarditis. (A) Uniform Manifold Approximation and Projection (UMAP) plot illustrating the clustering of neutrophil subtypes; (B) Distribution of neutrophil numbers across different phases of autoimmune myocarditis; (C) Dot plot demonstrating the expression of marker genes for distinct neutrophil types; (D) Pseudo-time trajectory analysis depicting the developmental trajectory of neutrophils; (E) Expression changes of marker genes during pseudo-time advancement, highlighting dynamic gene expression patterns throughout neutrophil development.

To understand the developmental trajectory of distinct neutrophil populations, we employed Monocle3 [26] for pseudo-time analysis. This revealed a highly ordered trajectory of neutrophil differentiation and maturation, starting with the Neutro3 cluster in the bone marrow and culminating in the Neutro2 cluster (Fig. 4C). Marker genes specific to each cluster were monitored for alterations during the trajectory. Rpl12 and Tuba1b had high expression levels at the beginning of the trajectory, but Oasl2 and Il18bp started with low levels, peaked in the middle, and then declined by the end of the trajectory. Prok2 and Mmp8 consistently showed increasing expression patterns toward the end of the journey (Fig. 4E).

### 2.5. Joint Learning of Conserved and Context-Specific Communication Patterns between Distinct Phases

Herein, we employed CellChat to investigate ligand-receptor interactions and decipher communication between immune cells across different phases of autoimmune myocarditis. By comparing the total number and strength of interactions, we observed a consistent increase from the normal to acute and subacute phases, followed by a reduction in the chronic myopathy phase (Fig. 5A). Notable alterations in signal transmission or reception were observed in various cell populations. Specifically, T cell monocyte complexes had heightened communication from normal to acute stages, whereas plasmacytoid dendritic cells had elevated considerable communication in the subacute phase but reduced communication in the chronic stage (Fig. 5B). Subsequently, we conducted a comprehensive analysis using joint manifold learning; categorizing inferred communication networks based on functional similarity. The pathways associated with these networks were plotted onto a shared two-dimensional surface and categorized into four separate groups (Fig. 5C). Group 1 consisted of pathways that influenced angiogenesis, extracellular matrix remodeling, cell adhesion, and B-cell activation. Group 2 was linked to pathways related to the innate immune response. Group 3 consisted of pathways associated with T-cell activation and function, whereas Group 4 comprised pathways crucial for proliferation. Furthermore, pathways linked to the activation and regulation of the immune system have been identified (Fig. 5D).

**Fig. 5.**
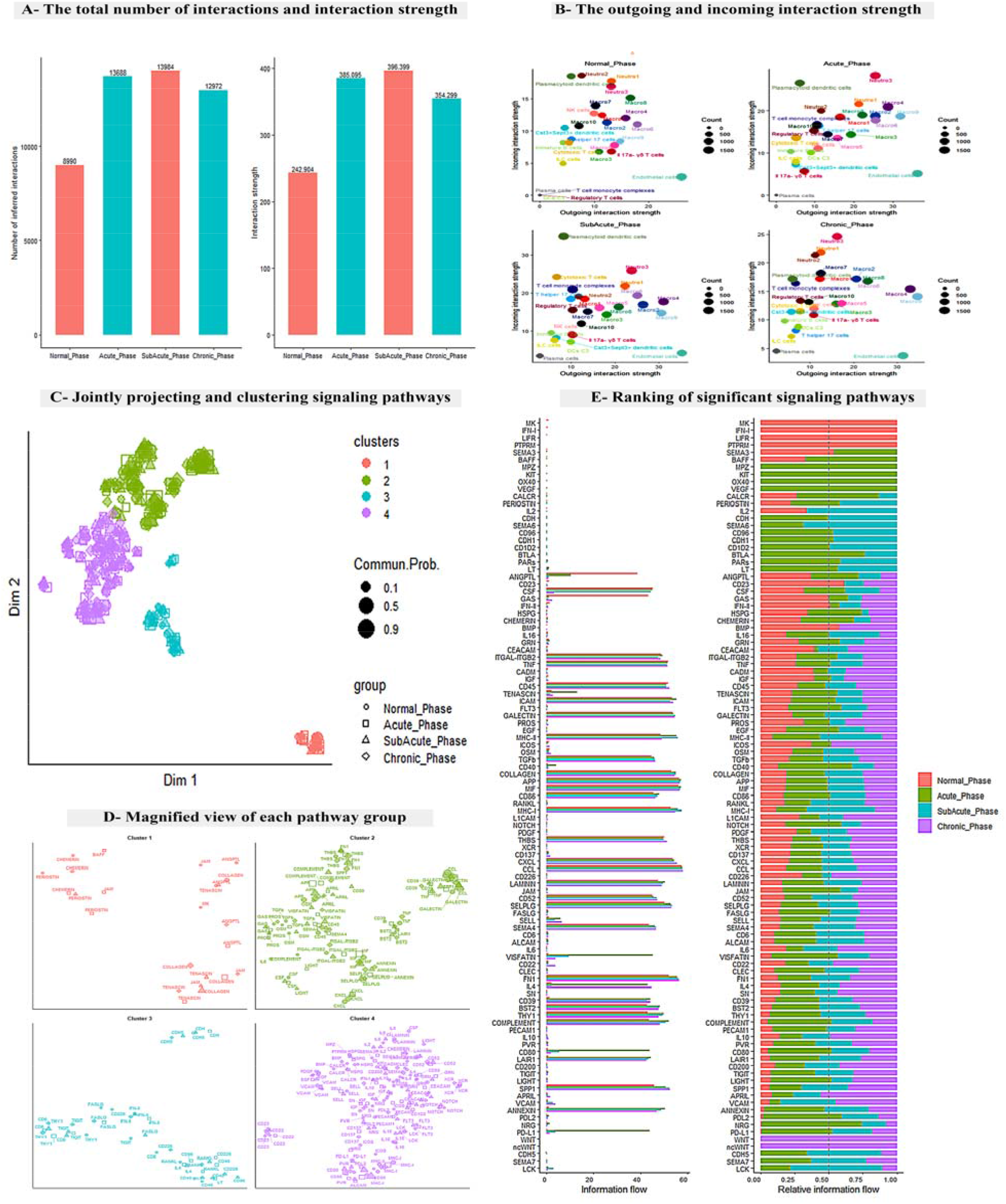
Joint Identification of Conserved and Context-Specific Communication Patterns between the Four Phases of Autoimmune Myocarditis. (A) Barplot illustrating the number of interactions and interaction strength across the different phases of autoimmune myocarditis; (B) Comparison of outgoing and incoming interaction strength visualized in 2D space; (C) Joint projection and clustering of signaling pathways, categorized into distinct groups; (D) Magnified view of each pathway group, providing detailed insights into the pathways’ characteristics; (E) Ranking of significant signaling pathways based on their activity and relevance across the four phases of autoimmune myocarditis.

We further evaluated information flow, representing the total likelihood of communication among the four datasets. Notably, 95 out of 105 circuits exhibited significant activity across normal, acute, subacute, and chronic myopathy phases (Fig. 5E). Specific pathways were found to be exclusively active during certain phases. For instance, pathways like MK, IFN-1, LIFR, and PTPRM were exclusively active during the normal phase, while pathways like MPZ, KIT, OX40, and VEGF were active only during the acute phase. Conversely, pathways like WNT and ncWNT were exclusively expressed during the chronic phase, associated with the proliferative process.

### 2.6. Comparison Analysis of Cell-Cell Communications between Normal and Acute Stages

To assess variations in the number and intensity of interactions between the normal and acute phases, we constructed a heatmap. The heatmap revealed an increase in the frequency of interactions between cytotoxic T cells, regulatory T cells, and macrophages, as well as heightened interactions between regulatory T cells and endothelial cells in addition to macrophages. Additionally, T cell monocyte complexes exhibited increased interactions with macrophages, neutrophils, and regulatory T cells. Similar trends were observed regarding the interaction strength, particularly between macrophages and endothelial cells, as well as regulatory T cells (Fig. 6A). We further investigated the similarities and differences in signaling pathways between the two phases by calculating the Euclidean distance between each pair of pathways shared in the two-dimensional manifold. Significant distances were observed for pathways such as BAFF, SPP1, and IL4, indicating distinct patterns of pathway activation between the normal and acute phases. Conversely, pathways like ANNEXIN, APRIL, PD-L1, PD-L2, PVR, and NRG showed relatively smaller distances, suggesting more conserved activation patterns (Fig. 6B).

**Fig. 6.**
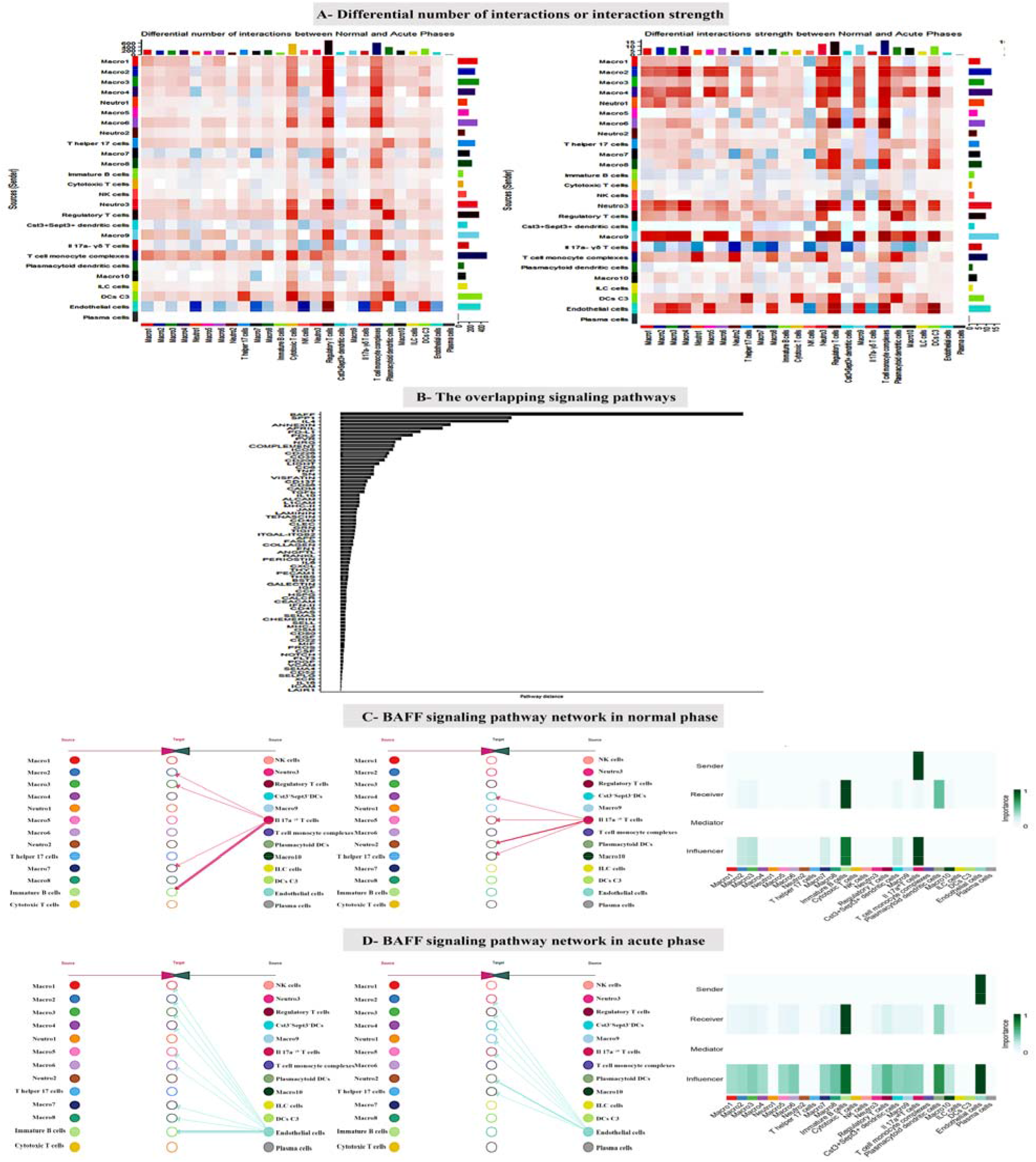
Comparison Analysis of Cell-Cell Communications between Normal and Acute Phases. (A) Heatmap illustrating several interactions and interaction strength between normal and acute phases; (B) Visualization of the overlapping signaling pathways between the normal and acute phases; (C) BAFF signaling pathway in the normal phase. The left figure represents the first 13 clusters, the median figure represents the last 13 clusters, and the right heatmap shows each cell group’s relative importance based on the BAFF signaling network’s four computed network centrality measures in normal phase; (D) BAFF signaling pathway in the acute phase. The left figure represents the first 13 clusters, the median figure represents the last 13 clusters, and the right heatmap shows each cell group’s relative importance based on the BAFF signaling network’s four computed network centrality measures in the acute phase.

A focused analysis was conducted on alterations in BAFF communications during the transition from the normal to the acute phase (Figs. 6C-D). During the normal phase, Il17a^-γδ^ cells were identified as the primary source of BAFF signaling. After careful examination, we have shown that Il17a^-γδ^ cells, with high sender scores, are the main sources of signaling for the BAFF pathway. Immature B cells and plasmacytoid dendritic cells have the most elevated receiver scores. Moreover, the analysis reveals that immature B cells and Il17a^-γδ^ cells exert the greatest influence on the information flow within a signaling network (Fig. 6C). During the acute phase, we identified the endothelial cells as the main origin of the BAFF signaling pathway. Our investigation found that endothelial cells are the primary origins of signaling in the BAFF pathway, exhibiting elevated sender scores. Immature B cells have the highest levels of receptor scores. Furthermore, when it comes to understanding the transmission of information in a signaling network, research has shown that the endothelium and immature B cells have the greatest impact on the flow of information pathways (Fig. 6D).

### 2.7. Comparison Analysis of Cell-Cell Communications between Acute, Subacute, and Chronic Phases

To assess variations in the number and strength of interactions between the acute, subacute, and chronic phases, we generated heatmaps. These heatmaps revealed a significant increase in the number of interactions between the T cell monocyte complex and most other cell types in the subacute phase when compared with the acute phase, which decreased during the chronic phase. Similarly, a similar interaction pattern was observed between Il17a^-γδ^ T cells and other cell types. Additionally, we noted a reduction in the connections between cytotoxic T cells and most other cell types in the subacute phase compared to the acute phase, with an increase observed during the chronic phase, particularly with macrophages, Il17a^-γδ^ T cells, and plasma cell clusters (Fig. 7A). Furthermore, based on our findings, there was an increase in the interaction strength between cytotoxic T cells and most cell types in the subacute phase. How-ever, this interaction strength decreased during chronic phases, including with plasmacytoid dendritic cells (Fig. 7A).

**Fig. 7.**
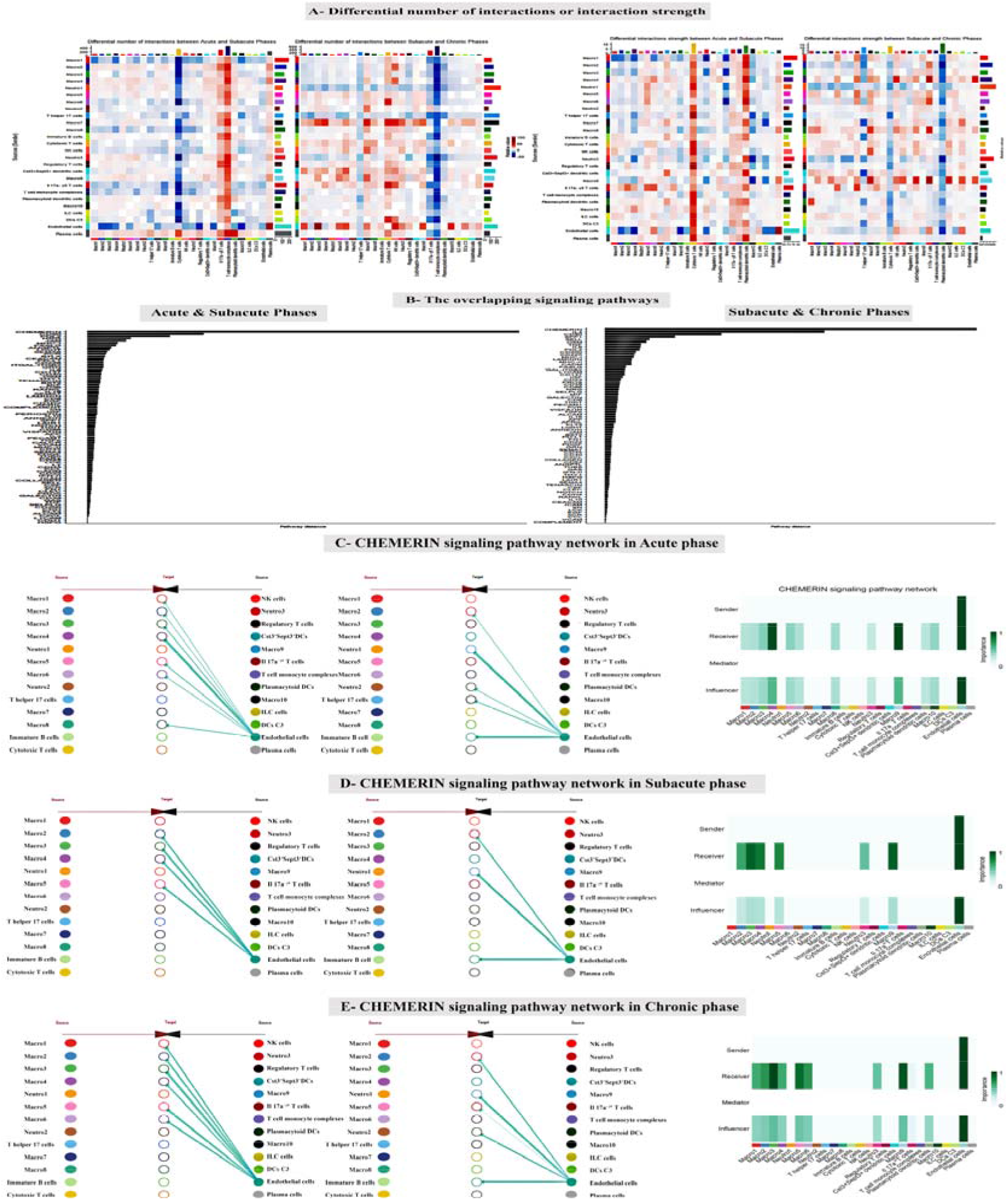
Comparison Analysis of Cell-Cell Communications between Acute, Subacute, and Chronic Phases. (A) Heatmaps illustrating several interactions and interaction strengths between the acute and subacute phases (left) and between subacute and chronic phases (right); (B) Visualization of the overlapping signaling pathways between acute and subacute phases (left) and between subacute and chronic phases (right); (C) CHEMERIN signaling pathway in the acute phase. The left figure represents the first 13 clusters, the median figure represents the last 13 clusters, and the right heatmap shows each cell group’s relative importance based on the CHEMERIN signaling network’s four computed network centrality measures; (D) CHEMERIN signaling pathway in the subacute phase; The left figure represents the first 13 clusters, the median figure represents the last 13 clusters, and the right heatmap shows each cell group’s relative importance based on the CHEMERIN signaling network’s four computed network centrality measures in subacute phase (E) CHEMERIN signaling pathway in the chronic phase, The left figure represents the first 13 clusters, the median figure represents the last 13 clusters, and the right heatmap shows each cell group’s relative importance based on the CHEMERIN signaling network’s four computed network centrality measures in chronic phase.

We conducted a focused analysis of the alterations in CHEMERIN communications throughout the three stages (Figs. 7C-D). In the acute phase, endothelial cells were the primary source of CHEMERIN signaling (Fig. 7C). After examination, we have shown that endothelial cells serve as the main sources of signaling in the CHEMERIN pathway, exhibiting elevated sources and sender scores. The acute phase has the greatest receiver scores in macro4, macro9, and endothelial cell clusters. Moreover, the analysis reveals that endothelial cells influence the information flow within a signaling network (Fig. 7C). In the subacute phase, we have also identified that the endothelial cells are the main origin of the CHEMERIN signaling pathway. Our investigation has shown that endothelial cells serve as the primary sources of signaling in the CHEMERIN pathway, exhibiting elevated sender scores. The receiver scores are highest in the clusters of endothelial cells, macro2, macro3, macro4, macro5, and macro9. Furthermore, when it comes to understanding the transmission of information in a signaling network, research has shown that endothelial cells have the greatest impact on the flow of information (Fig. 7D). In the chronic phase, similar findings were observed, with the addition of macro1, macro6, clusters, and plasmacytoid dendritic cells showing significant receiver scores and influencer scores (Fig. 7E).

## 3. Discussion

Single-cell technologies have emerged as powerful tools for analyzing and distinguishing between various cell types within both healthy and diseased tissues. Recent applications of these techniques have provided valuable insights into the cellular composition of the normal human heart [27,28].

Myocarditis involves a sophisticated inflammatory reaction in the heart resulting from abnormal immunological mechanisms, which can be observed clinically and histologically [29]. The condition usually advances through three specific stages: acute inflammation, subacute inflammation, and chronic myopathy. Managing myocarditis is challenging due to the need to target autoimmune responses or decrease immune cell activity involved in inflammatory cardiomyopathy without causing harm to the host [30]. The comprehensive analysis of ligand-receptor interactions using CellChat has unveiled intricate patterns in cell-cell communication across multiple stages of myopathy, including normal, acute, subacute, and chronic phases. The interactions among different subsets of immune cells reflect the dynamic nature of immune responses during distinct periods. Notably, our research has identified distinct cell populations playing pivotal roles in the altered communication dynamics. During the acute phase, levels of T cell-monocyte complexes were notably elevated, whereas plasmacytoid dendritic cells exhibited a significant increase during the subacute phase, followed by a decline in the chronic stage.

Employing joint manifold learning and classification, we categorized the predicted communication networks into four distinct pathway groups. Group 1 encompassed pathways associated with critical factors such as angiogenesis, extracellular matrix remodeling, cell adhesion, and B-cell activation. Group 2 plays a crucial role in the body’s natural defense systems, while Group 3 is associated with pathways closely related to T cell activation and function. Group 4 pinpointed pathways essential for cell growth. An analysis of information flow across datasets unveiled intriguing insights. Most routes (95 out of 105) exhibited significant activity across all stages. However, specific pathways demonstrated phase-specific activity, with four pathways exclusively active during the normal phase, four during the acute phase, and two during the chronic phase. The complex results highlight the varied involvement of immune cells at different points in myopathy, offering useful guidance for focused research and treatment approaches. Recent studies have focused on examining immune cells to better understand autoimmune myocarditis. Eriksson et al.[31] established an experimental model of myosin-induced autoimmune myocarditis by utilizing dendritic cells that were loaded with myosin peptides. The activation of the T cell system is regarded as the primary pathophysiological mechanism underlying autoimmune myocarditis and autoimmune inflammatory cardiomyopathy[32]. Specifically, mice lacking the T-box transcription factor TBX21 (T-bet), critical for TH1 cell development and IFNγ generation, are more susceptible to autoimmune myocarditis due to heightened IL-17 production[33]. Antigen-presenting cells release IL-23, which promotes the production of TH17 cells, resulting in an elevated TH17/Treg cell ratio and inducing autoimmune myocarditis in rats.

Monocytes and macrophages represent the primary types of inflammatory cells observed in both human and experimental myocarditis cases[34]. The increased detection of T cell-monocyte complexes during the acute and subacute phases underscores their potential role in autoimmune myocarditis progression. Gene ontology and pathway analyses offer crucial insights into the functional implications of these complexes, emphasizing their involvement in signal transduction, immune response regulation, T cell proliferation, and activation. Moreover, their involvement in key pathways such as T cell receptor signaling, MAPK signaling, and leukocyte transendothelial migration highlights their importance in coordinating adaptive immune responses and inflammation. Furthermore, it was observed that immune cells exhibited a significant level of contact with most cells, except during the chronic phase, where the number of connections was reduced. Specifically, macrophages emerged as the most prevalent population. Our research identified 10 separate sub-clusters of macrophages, each demonstrating distinct dynamics. Trajectory analysis was employed to elucidate the process of monocyte/macrophage infiltration into the infarcted region. Initiating from Ly6c2^hi^ monocytes, cells transitioned from monocytes derived from macrophages to clusters involved in antigen presentation, characterized by an oxidative stress pathway. Subsequently, the complement and phagocytosis macrophage pathway emerged. This trajectory analysis was corroborated by gene expression profiles across pseudo-times, highlighting dynamic alterations. Initially, genes related to macrophage-derived monocytes exhibited high expression levels but decreased over time. In contrast, genes associated with antigen presentation and the processes of complement and phagocytosis reached their peak levels during the pseudo-time of monocytes produced from macrophages. Notably, while the levels of H2-Eb1 markers decreased, C1qc indicators continued to rise. The expression of the oxidative stress marker Ltc4s remained consistently high, whereas the inflammatory marker Treml4 exhibited a regular expression pattern with occasional deviations throughout the pseudo-time of oxidative stress in macrophages.

In summary, our research provides valuable insights into the diversity and changes of macrophage populations over time in relation to immunological responses in the heart [34]. Furthermore, we identified that macrophages are essential in acquiring CHEMERIN together with endothelial cells throughout the shift from the acute phase to the chronic phase.

Neutrophils play a crucial role in sustaining inflammation through a distinct mechanism called NETosis, wherein neutrophil extracellular traps are formed. Research has demonstrated the involvement of NETosis in promoting cardiac inflammation in mice with experimental autoimmune myocarditis (EAM) [35]. Furthermore, there is a significant association between the amount of neutrophils present in the heart and the degree of acute myocardial inflammation in these animals [36]. In this study, three distinct groups of neutrophils were identified. The quantity of neutrophils was notably higher during the acute and subacute phases compared to the normal group. The Neutro3 cluster, primarily characterized by the expression of Tuba1b and Rpl12, was found to originate from the bone marrow and undergo differentiation inside the BM. Functional investigation of the Neutro2 and Neutro3 clusters revealed their involvement in neutrophil degranulation, leukocyte movement, and the structure of the actin cytoskeleton. On the other hand, the Neutro1 cluster displayed a higher level of expression of Oasl2 and Il18bp, which is indicative of its involvement in the defensive response, neutrophil degranulation, and cytokine production. In addition, the study discovered a rather well-organized pattern of neutrophil populations, with the Rpl12 and Tuba1b genes exhibiting high levels of activity at the beginning of the study. Conversely, the activity of the Oasl2 and Il18bp genes was lower at the beginning of the experiment, but it grew in the middle of the experiment, and it declined as the experiment progressed. Particularly noteworthy is the fact that the Prok2 and Mmp8 genes exhibited consistent expression patterns, with a discernible increase seen toward the conclusion of the trajectory. Further examination of these distinct types of neutrophils may lead to the development of specific treatment approaches for myocarditis, offering promising avenues for targeted intervention.

## 4. Methods

### 4.1. Data Collection

Raw single-cell RNA sequencing files were obtained from the Gene Expression Omnibus (GEO), specifically from the dataset GSE142564 [25].

### 4.2. Single-cell RNA Sequencing Analysis and Identification of Marker Genes

The gene expression data for each sample was meticulously processed in the R environment and converted into Seurat objects using the Seurat R tool. A rigorous quality assurance method was established to guarantee the accuracy and dependability of the data. Cells with over 40,000 unique molecular identifiers (UMIs), those with more than 5,000 genes expressed or less than 200 expressed genes were excluded. Normalization and variance stabilization were performed using the sctransform technique developed in Seurat. To mitigate the influence of cell cycle effects, comprehensive regression analysis was conducted as outlined in the accompanying vignette [https://satijalab.org/seurat/v3.0/cell_cycle_vignette.html].

Integration of datasets from all four conditions was completed. Subsequently dimensionality reduction was performed using Principal Component Analysis (PCA) and Uniform Manifold Approximation and Projection (UMAP) in 40 dimensions [37]. UMAP visualization in Seurat identified 26 distinct clusters at a resolution level of 0.7. Marker genes inside cell clusters were identified by comparing cells in a single cluster to those in all other clusters using Seurat’s “FindConservedMarkers” function. Following that, genes with high expression in the T cell monocyte complex were subjected to gene ontology and KEGG pathway analysis. The data was visualized using the ggplo2 package.

### 4.3. Cell Trajectory and Pseudotime Analysis

Trajectory inference and pseudotime modeling were conducted using Monocle 3 (v1.3.4) [38]. The “learn_graph” function was utilized to systematically rank cells based on pseudotime and identify their trajectories. Visualization of trajectory dynamics was achieved using the “plot_cells” function.

### 4.4. Cell Communication Analysis

CellChat was used according to recognized methods to understand the complex cell signaling and communication networks involved in different stages of autoimmune myocarditis [39]. An extensive analysis of known ligand-receptor interactions among various immune cell types was carried out using the entire CellChat database, revealing possible signaling pathways among certain immune cell subgroups.

## 5. Conclusion

Overall, our use of single-cell RNA sequencing has provided a detailed and ever-changing description of the immune system’s characteristics in autoimmune myocarditis. Through the process of delineating and categorizing different immune cell populations, such as T cell-monocyte complexes, macrophages, and neutrophils, we have clarified their unique functions and changes throughout time, emphasizing their crucial roles in the advancement of diseases.

By conducting a thorough analysis of the communication networks among cells, researchers have discovered both persistent and phase-specific patterns. These patterns have yielded vital observations regarding the evolving characteristics of immune responses during the different phases of disease therapy. These insights have improved our comprehension of autoimmune myo-carditis at the cellular level and have also shown potential routes for the development of customized therapeutic treatments.

This comprehensive examination of cellular and molecular dynamics has laid a strong groundwork for future research and advancements in precision medicine approaches that explicitly target the effective treatment of autoimmune myocarditis.

## 6. Limitations and Future Perspectives

Our study acknowledges certain limitations of single-cell RNA sequencing in providing insights into the immunological landscape of autoimmune myocarditis. Firstly, the resolution of the technology may not adequately catch rare cell populations, which could result in underrepresentation or misinterpretation of specific cell types. In addition, the sample size in our study may have been restricted, which could have impacted the applicability of our findings. Moreover, the analysis of single-cell RNA sequencing data primarily depends on computational methods and bioinformatics tools, which can potentially induce biases or inaccuracies. Hence, it is imperative to conduct additional experimental verification of the main discoveries in order to guarantee the dependability of our results. In addition, our study provides insight into the time-related changes in immune responses in auto-immune myocarditis. However, to fully understand the specific timing of cellular interactions and signaling pathways, further research employing longitudinal studies is necessary.

Our research uncovers multiple perspectives when examining the future. Integrating data from single-cell RNA sequencing with other omics approaches, like as proteomics and metabolomics, can lead to a comprehensive understanding of the molecular mechanisms underlying autoimmune myocarditis. Moreover, it is imperative to conduct further experimental validation of crucial discoveries, including functional assays and in vivo models, in order to ascertain the roles of specific cell types and pathways identified in our research. In order to use our findings in a clinically meaningful manner, it is necessary to conduct thorough validation in patient cohorts and create targeted therapeutic approaches tailored to specific disease subtypes. Longitudinal studies that track immune cell dynamics and molecular profiles over time in patients with autoimmune myocarditis can be used to identify new biomarkers for early diagnosis and prognosis. Furthermore, these examinations can provide valuable insights into the progression of disease. Ultimately, our research findings establish the basis for identifying potential pharmaceutical targets and therapeutic strategies for treating patients with autoimmune myocarditis. Future research should prioritize the screening and development of new pharmacological medicines that can modify dysregulated immune responses and restore cardiac homeostasis.

## Institutional Review Board Statement

Not applicable. Informed Consent Statement: Not applicable.

## Data Availability Statement

Access to the data is available in the Gene Expression Omnibus (GEO) database under accession number [GSE142564].

## Acknowledgments

The authors extend their appreciation to the Deanship of Scientific Research at King Khalid University for funding this work through a large group Research Project under grant number RGP2/17/44. The author extends her appreciation to the Deanship of Scientific Research at Princess Nourah bint Abdulrahman University Researchers Supporting Project number (PNURSP2024R465), Princess Nourah bint Abdulrahman University, Riyadh, Saudi Arabia. Furthermore, we sincerely appreciate Edigenomix Scientific Co., Ltd. for their expert editing and proofreading, which greatly improved the clarity and quality of our manuscript. Their meticulous attention to detail and support in refining the document for publication is gratefully acknowledged.

